# Modeling the Effects of Propofol on Brain Activity: Insights from Computational EEG Studies

**DOI:** 10.1101/2024.10.06.616892

**Authors:** Saeed Zahran

## Abstract

The effects of anesthetic agents like propofol on brain activity provide critical insights into how consciousness is altered during anesthesia. This article presents a computational approach to modeling the impact of propofol on the brain’s electrical activity using the COALIA model, a large-scale neural mass model of human EEG designed for consciousness research. Propofol enhances GABA_A-mediated synaptic inhibition, leading to characteristic EEG signatures such as alpha oscillations (8-12 Hz) and slow-wave oscillations (0.1-1.5 Hz). We model these effects by modulating the parameters of postsynaptic potentials and synaptic strength, simulating the action of propofol at both the molecular and network levels. COALIA’s detailed representation of thalamo-cortical and cortico-cortical networks provides realistic simulations of brain dynamics under anesthesia, closely replicating experimental EEG data. Our findings highlight the importance of network-level interactions in producing the distinctive EEG changes seen during propofol-induced unconsciousness. This research paves the way for future applications in personalized anesthesia monitoring and consciousness assessment through EEG modeling.

## I. Introduction

Understanding the effects of anesthetic agents, particularly propofol, on brain activity has profound implications for both neuroscience and clinical applications. Propofol, a widely used anesthetic, is known to cause a reversible loss of consciousness by modulating neural circuits at both molecular and network levels. Despite its routine use, the precise mechanisms by which propofol alters consciousness remain incompletely understood. This gap in understanding is largely due to the complex interplay between molecular actions, such as its potentiation of GABA_A receptors, and macroscopic changes in brain network activity, as observed through electroencephalographic (EEG) recordings (1, 2).

General anesthesia produces notable changes in brain dynamics, including distinct shifts in EEG signals. Propofol, for example, induces specific spectral properties such as alpha oscillations (8–12 Hz) and Slow Wave Oscillations (SWO, 0.1–1.5 Hz), which are characteristic of the anesthetic state (3). At the molecular level, propofol enhances synaptic inhibition by potentiating the activity of GABA_A receptors—the brain’s primary inhibitory neurotransmitter— prolonging the inhibitory postsynaptic potential (IPSP) duration (4, 5). This prolongation reduces neuronal excitability and plays a critical role in the global suppression of brain activity associated with unconsciousness. However, while these molecular effects are well-established, less is known about how they translate to the large-scale network dynamics observed in the EEG.

Recent research shows that the actions of propofol on GABA_A receptors lead to widespread alterations in brain rhythms, particularly in slow-wave and alpha bands, reflecting a reorganization of cortico-cortical and thalamo-cortical connectivity (6, 7). The thalamo-cortical system, crucial for sustaining consciousness, is significantly affected by anesthetic agents like propofol, which disrupts normal connectivity patterns, contributing to the breakdown of network-level information processing that underlies unconsciousness (8). While neural mass models (NMM) and thalamo-cortical neural models have been applied to simulate brain rhythms during various conscious states, including sleep and anesthesia (9, 10), there is a need for more integrated models that capture the link between molecular actions and brain-wide EEG changes.

The present study utilizes the COALIA model, a computational tool designed to simulate brain-scale electrophysiological activity under different states of consciousness (11). COALIA incorporates both cellular-scale connectivity patterns and large-scale anatomical connections, providing a framework for simulating how molecular-level interactions, such as the effects of propofol on GABAergic transmission, manifest as changes in EEG rhythms. This approach bridges the gap between molecular mechanisms and macroscopic brain dynamics, offering insights into how anesthetic-induced alterations in synaptic activity lead to the characteristic EEG signatures of unconsciousness, including the co-occurrence of delta and alpha oscillations (12, 13).

By simulating the effects of propofol, COALIA allows us to explore the disruption of brain network dynamics during anesthesia. Specifically, propofol-induced unconsciousness is characterized by a breakdown of cortico-cortical and thalamo-cortical communication, which is consistent with prominent theories of consciousness. The global workspace theory, for instance, suggests that consciousness relies on the functional integration of distributed cortical regions (14), while the integrated information theory (IIT) highlights the importance of both integration and differentiation of information across brain networks (15). In the context of these theories, the ability of COALIA to simulate network dynamics under anesthesia provides a powerful tool for investigating the neural correlates of consciousness and the mechanisms by which anesthetics such as propofol alter brain function.

In this study, we aim to connect the molecular and network levels of brain activity by using COALIA to simulate the effects of propofol on GABAergic transmission and large-scale brain connectivity. Through these simulations, we investigate how the molecular effects of propofol lead to characteristic EEG oscillations and disrupt information processing across brain networks, ultimately resulting in the loss of consciousness. Our findings contribute to the growing body of research focused on understanding the neural basis of consciousness and the role of anesthetics in modulating brain dynamics (16, 17).

## II. Material and methods

### Computational Model: COALIA Framework

The COALIA model is a large-scale computational framework designed to simulate neuronal activity patterns from local microcircuits to whole-brain dynamics, replicating brain activity during various states of consciousness, such as wakefulness, sleep, and anesthesia. Each computational unit, called a neural mass, represents the activity of local populations of neurons, including pyramidal cells and GABAergic interneurons, which generate local field potentials (LFPs) and EEG signals. These neural masses are spatially distributed across 66 cortical regions based on the Desikan anatomical parcellation (20) and connected via long-range cortico-cortical and thalamo-cortical projections. The structural connectivity is derived from Diffusion Tensor Imaging (DTI) data (21). Conduction delays are incorporated to account for the finite propagation time of neural signals across these regions.

### Propofol Mechanisms and Modeling

Propofol, an anesthetic agent, enhances GABAergic transmission by modulating GABA_A receptors, leading to a prolongation of inhibitory postsynaptic potentials (IPSPs) and reducing neural excitability (1). The COALIA model represents these effects using a dimensionless scale factor (λ) that reflects the concentration of propofol. The following equations describe the effects of propofol on synaptic transmission:

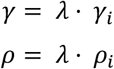

Here, γ_*i*_ and ρ_*i*_ represent the baseline values of the inhibitory rate constant and synaptic strength, respectively, while **λ** increases in proportion to propofol concentration (2, 3).

### Model Parameters with Propofol

Key parameters related to post-synaptic potentials (PSPs) are scaled according to propofol concentration, as shown below:

**Table 1:** Model parameters interpretation and values.

| Parameter | Significance | Initial Value | With Propofol |
| --- | --- | --- | --- |
| B (mV) | Inhibitory PSP amplitude (dendritic targeting interneurons) | 30 | 84 |
| G (mV) | Inhibitory PSP amplitude (somatic targeting interneurons) | 20 | 57 |
| b (ms) | PSP rate constant (dendritic targeting interneurons) | 104.87 | 314.61 |
| g (ms) | PSP rate constant (somatic targeting interneurons) | 20.97 | 43.69 |

These changes reflect the increased IPSP duration and reduced neural excitability due to propofol’s effects on GABA_A receptors (4, 5).

### Thalamocortical Networks

In the COALIA model, we considered a thalamic population and 66 cortical populations to simulate realistic activity across the neocortex during anesthesia (6). Each population’s time course represents the activity of a cortical region, derived from Desikan’s anatomical parcellation (20). Structural connectivity, based on DTI data, includes spatial distance-based propagation delays between regions. Propofol-induced unconsciousness is marked by coherent alpha rhythms (9–13 Hz) in the frontal cortices (8). The reciprocal coupling between the thalamus and cortex generates sustained alpha activity, which synchronizes across cortical regions through the thalamus, resulting in high spatial coherence (9)

### EEG Effects of Propofol

Propofol anesthesia is associated with significant changes in EEG rhythms, including beta oscillations (13–25 Hz) observed before the loss of consciousness, and slow-wave (0.1–1 Hz), delta (1–4 Hz), and alpha (∼10 Hz) oscillations during unconsciousness (10, 11). The COALIA model simulates these changes, focusing on the frontal cortical regions, where propofol’s effects on neural connectivity are most pronounced.

### Simulation Protocol

Simulations were conducted using the COALIA framework, adjusting parameters to reflect varying levels of thalamo-cortical and cortico-cortical connectivity at different concentrations of propofol. The simulations aimed to replicate delta and alpha oscillations, which are critical indicators of propofol-induced unconsciousness (12).

The simulated EEG data were analyzed using spectral decomposition techniques, focusing on power changes in the delta (0.1–4 Hz) and alpha (8–12 Hz) bands, which are strongly associated with propofol’s effects on brain activity.

### Forward Model: From Cortical Activity to EEG

After simulating local field potentials (LFPs) in the 66 cortical regions, the forward model was applied to generate realistic scalp EEG signals. This process involved solving the forward problem using the Boundary Element Method (BEM) (13) and reconstructing the EEG data from the simulated cortical activity.

### Head Model Construction

A realistic head model was built using the Colin 27 template MRI (14) in Brainstorm (15). The head model included three nested surfaces representing the cortex, skull, and scalp, with conductivity values of 0.33 S/m, 0.008 S/m, and 0.33 S/m, respectively (16).

### Leadfield Matrix

The neural activity was projected onto the scalp EEG electrodes using a leadfield matrix (A). The matrix was calculated for 257 high-density EEG electrode positions, and a reduced 66 × 257 matrix (G) was generated by summing the contributions from each cortical region (17).

### EEG Signal Generation

The spatio-temporal cortical activity (S) was projected onto the scalp using the leadfield matrix to generate the simulated EEG signals (X) according to the equation:

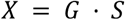

This method generated realistic EEG signals that were then analyzed to capture the characteristic effects of propofol on brain activity.

### Data Analysis

The simulated EEG data were subjected to time-frequency analysis to extract power spectra in the delta (0.1–4 Hz) and alpha (8–12 Hz) frequency bands.

The accuracy of the COALIA model was validated by comparing the simulated EEG data to experimental EEG recordings from patients undergoing propofol-induced anesthesia. Spectral power distributions and functional connectivity metrics were used to verify that the model accurately reproduced the neural dynamics associated with unconsciousness (19).

By incorporating propofol’s molecular mechanisms into a large-scale brain model, the COALIA framework provides a detailed understanding of how anesthetic agents affect brain dynamics, from molecular interactions to network-level activity, as captured in EEG data during anesthesia.

## III. RESULTS

The COALIA model was used to simulate the effects of propofol on brain activity, and the results were analyzed through comparisons between simulated and real EEG signals during anesthesia. Below is a summary of the key results:

### EEG Effects of Propofol

Figure 1A shows a time-frequency spectrogram of the EEG data obtained from a healthy volunteer who underwent propofol-induced general anesthesia. In the spectrogram, we observe the emergence of prominent delta (0.1–4 Hz) and alpha (8–12 Hz) oscillations as the patient transitions into unconsciousness. The data also highlight how the characteristic slow-wave and alpha rhythms dominate as anesthesia deepens. Figure 1B presents 10-second raw EEG data from the same subject, while Figure 1C demonstrates the bandpass-filtered signals, clearly isolating the slow and alpha oscillatory components. These results illustrate the spectral features of propofol anesthesia, supporting the suppression of high-frequency beta oscillations and the enhancement of slow and alpha rhythms.

**Figure 1:**
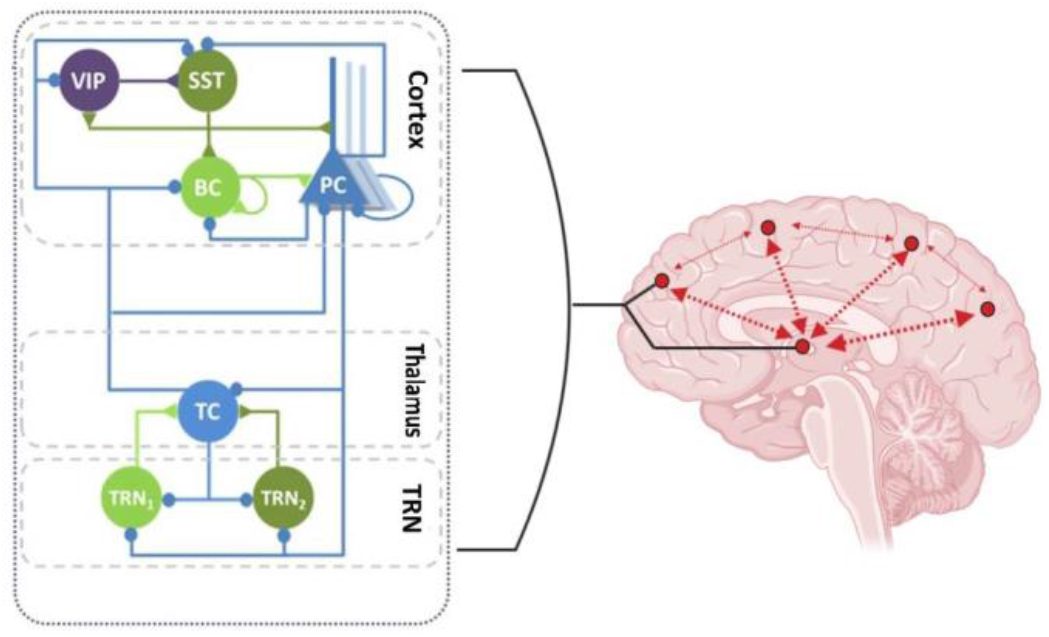
Thalamocortical system: in high thalamocortical connectivity state (i.e. low cortico-cortical connectivity)

**Figure 1:**
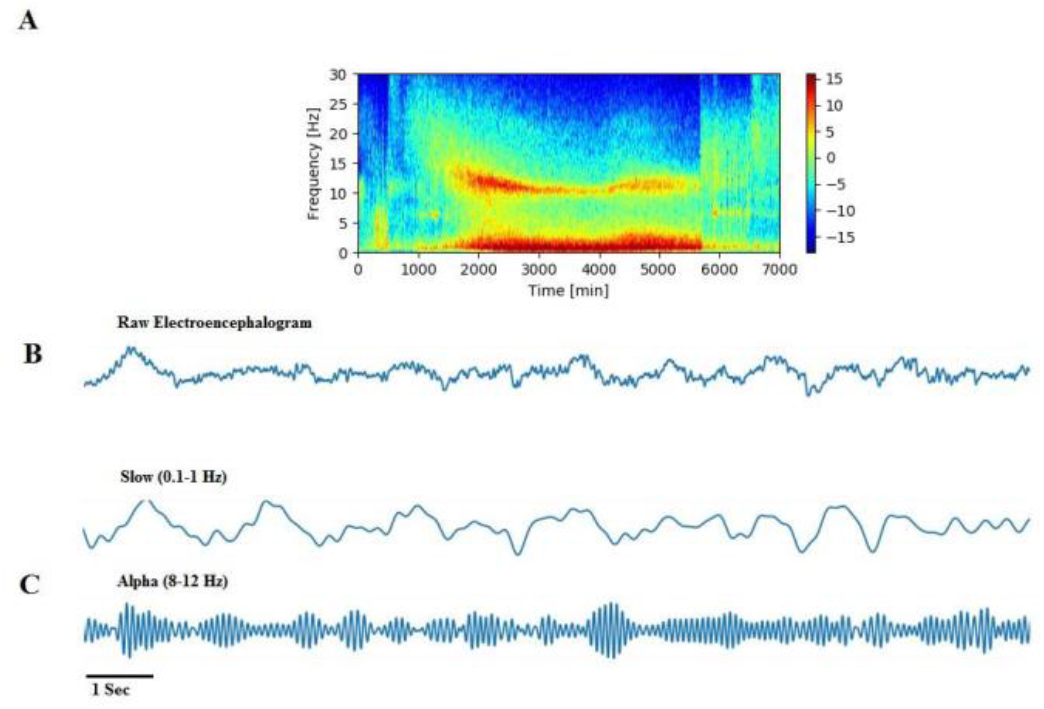
Representative individual spectrogram (time-frequency domain) and time-domain EEG data obtained during propofol-induced unconsciousness [8]. A) Spectrogram of a healthy volunteer who received propofol general anesthesia. (B) Representative 10 seconds of electroencephalogram data from A. (C) Bandpass-filtered EEG signals from the raw tracings to emphasize the EEG frequency content.

### Simulation of Post-Synaptic Response Under Propofol

Figure 2 shows the post-synaptic response function for excitatory and inhibitory neurons under the effect of propofol. The light curve represents excitatory post-synaptic potentials (EPSPs), while the bold solid curve represents inhibitory post-synaptic potentials (IPSPs) at a baseline condition (λ = 1), and the dashed line depicts the IPSPs under the influence of propofol (λ = 1.5). The presence of propofol significantly prolongs the duration of the IPSP, demonstrating how the anesthetic enhances GABAergic transmission. This prolongation reduces neuronal excitability, contributing to the global slowing of brain rhythms, a hallmark of propofol anesthesia.

**Figure 2:**
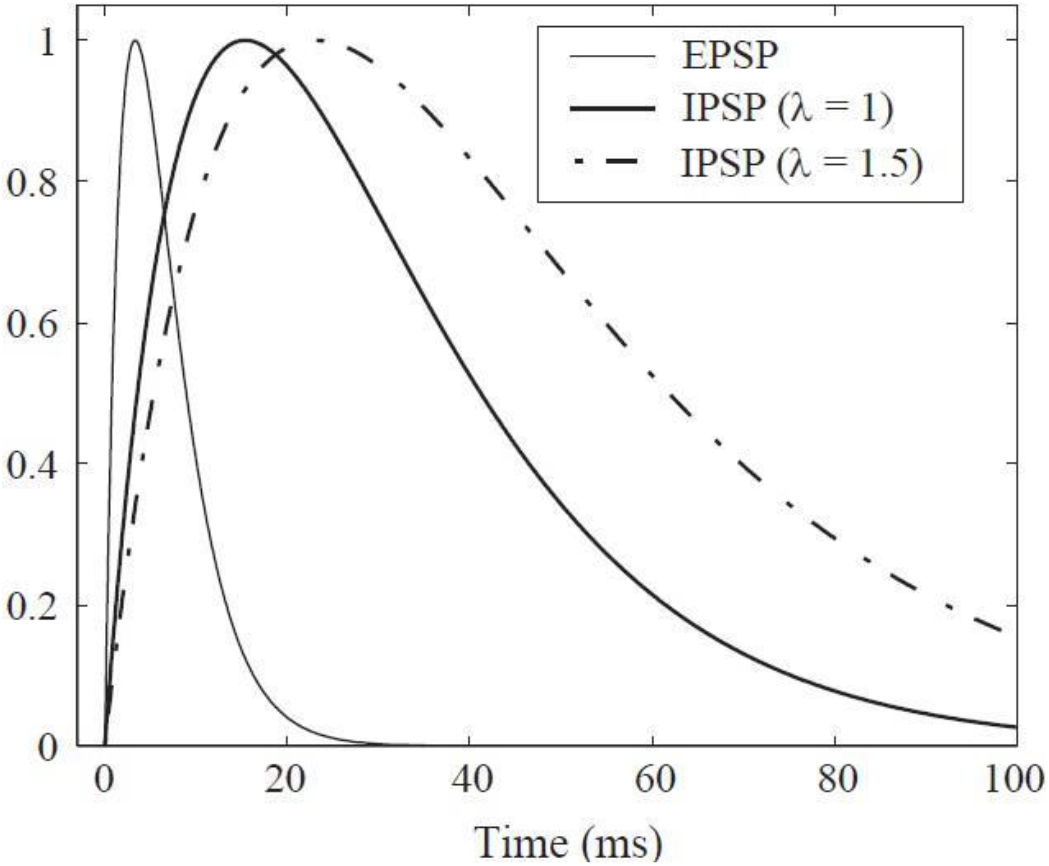
Post-synaptic response function for postsynaptic neurons that are excitatory (solid curve, EPSP), inhibitory at baseline (bold solid, IPSP λ = 1), and anesthetically altered under propofol (bold dashed, IPSP λ = 1.5). The λ symbol represents the dimensionless anesthetic-effect scale factor that increases the duration of IPSPs under the influence of propofol.

### Simulated vs. Real EEG Activity Under Propofol

Figure 3A compares the simulated EEG signals generated by the COALIA model with real EEG recordings from a patient under propofol anesthesia. The model successfully replicates the observed EEG features, such as the slow-wave and alpha oscillations that dominate during unconsciousness. The power distribution across frequency bands for both the simulated and real EEG signals, shown in Figure 3B, further validates the accuracy of the simulation. The delta band dominates the power spectrum in both the real and simulated EEG, consistent with the expected effects of propofol.

**Figure 3:**
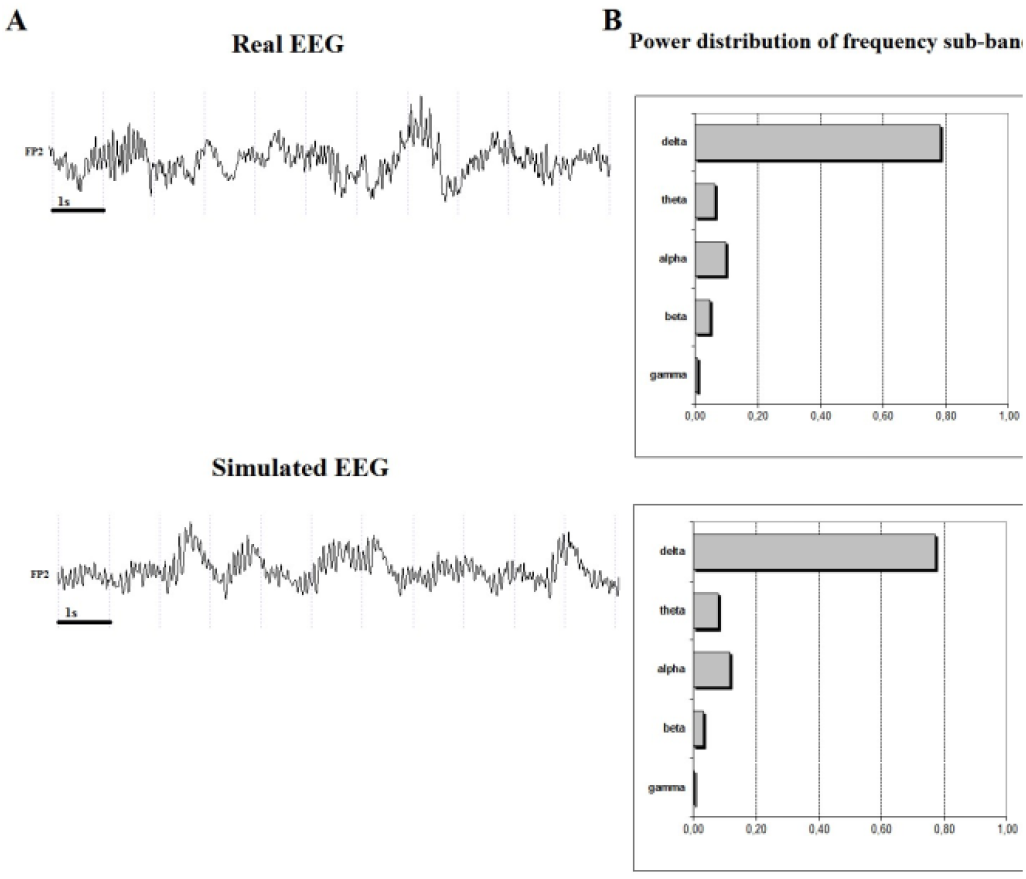
Comparison of simulated versus experimental effects of propofol on EEG activity. A) Simulated versus real EEG signals during propofol anaesthesia. Signals simulated with the whole-brain model (COALIA) using a high level of thalamo-cortical connectivity. B) The power distribution of the frequency bands in simulated signals is highly similar to the one in scalp EEGs recorded from human subjects during propofol anaesthesia.

### Comparison of Real and Simulated EEG Across Cortical Regions

Figure 4 presents a direct comparison between real EEG recordings and simulated EEG signals generated using the COALIA model. The simulated signals, obtained from 66 cortical regions, closely match the real EEG data in terms of both temporal dynamics and spatial distribution. The synchronization across cortical regions, mediated by thalamo-cortical interactions, is evident in both real and simulated data, further supporting the model’s ability to capture the large-scale brain dynamics induced by propofol.

**Figure 4:**
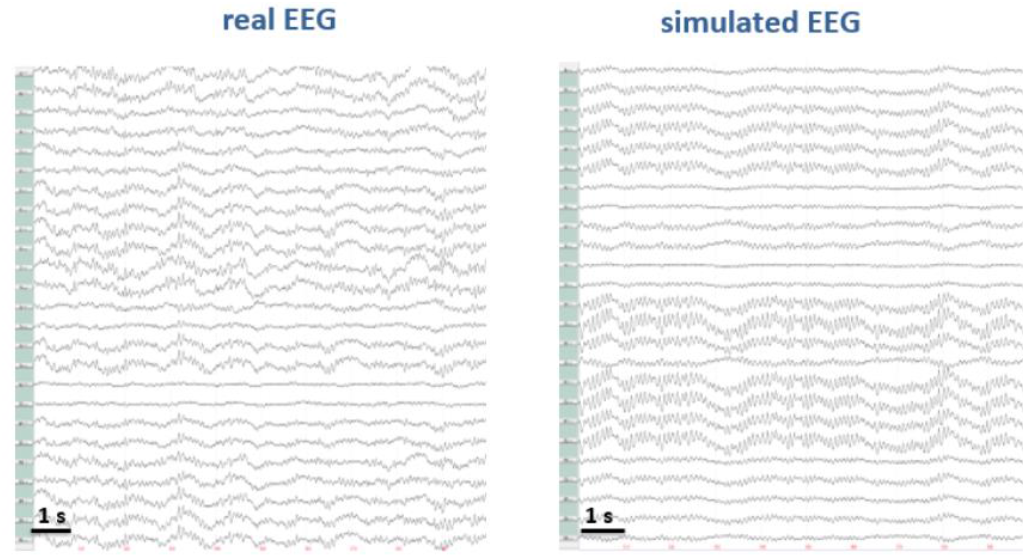
Comparison of real versus simulated EEG signals under propofol-induced anesthesia. Left panel shows the real EEG signals recorded from a human subject during propofol anesthesia. Right panel displays the corresponding simulated EEG signals generated using the COALIA model, reflecting similar oscillatory patterns as seen in real EEG, particularly in the slow-wave and alpha frequency bands. Both EEG traces highlight the capability of the model to replicate the characteristic brain rhythms observed during anesthesia.

### Evolution of GABA_A Decay Time and its Effect on EEG

Figure 5 illustrates how changes in the decay time of GABA_A receptors, modulated by propofol concentration, influence EEG activity. As the decay time increases, the slow-wave activity (SWA) power in the EEG increases, reflecting the enhanced inhibitory effects of propofol. This figure shows the evolution of GABA_A receptor decay time (top panel), the corresponding amplitude of the EEG signal (middle panel), the time-frequency representation of the EEG (third panel), and the increase in SWA power (bottom panel) with higher propofol concentrations.

**Figure 5:**
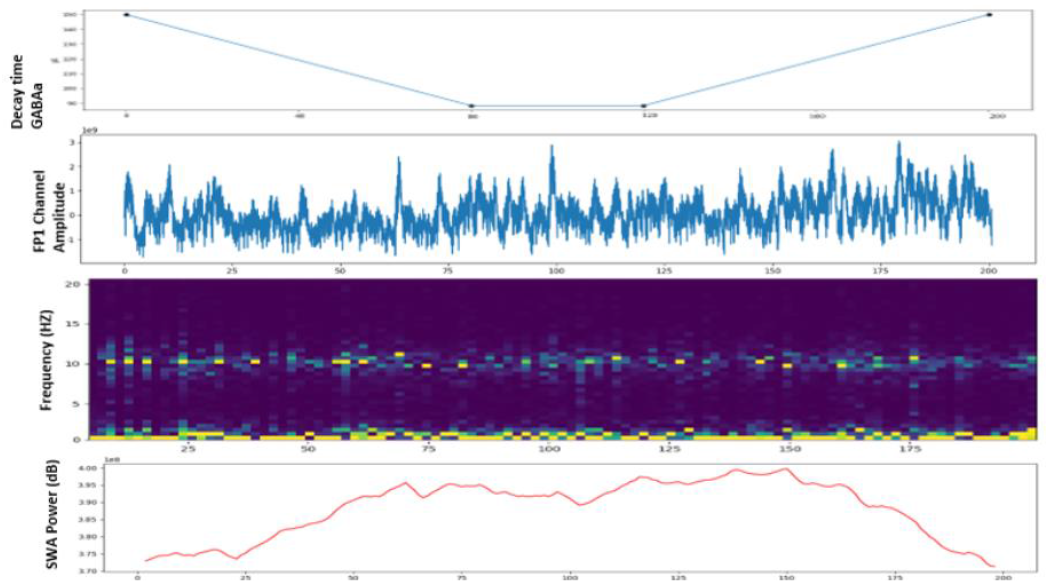
Evolution of GABAA_AA decay time and its impact on EEG features. The top panel shows the changes in GABAA_AA decay time, simulating the effect of increasing propofol concentration. The second panel presents the corresponding EEG amplitude recorded from the FP1 channel. The third panel shows the spectrogram of EEG signals across various frequencies, with notable changes in low-frequency activity. The bottom panel illustrates the slow-wave activity (SWA) power (0.1–1 Hz), which increases as GABAA_AA decay time lengthens, before decreasing toward the end of the simulation. This figure highlights the relationship between GABAA_AA modulation and the resulting dynamics in slow-wave oscillations.

### Interpretation of Results

These results demonstrate the COALIA model’s capacity to simulate the neural dynamics associated with propofol-induced anesthesia. The prolonged inhibitory effects, mediated by GABA_A receptors, lead to the suppression of high-frequency activity and the emergence of slow-wave and alpha rhythms. The comparison between simulated and real EEG data confirms that the model accurately captures the key electrophysiological changes observed during anesthesia.

By integrating thalamo-cortical interactions, the COALIA model provides a mechanistic understanding of how propofol alters consciousness, transitioning the brain into a state dominated by large-scale synchrony and slow oscillatory activity. These findings offer valuable insights into the neural correlates of unconsciousness and establish the utility of computational modeling for exploring the effects of anesthetic agents on brain function.

## IV. DISCUSSION AND CONCLUSION

The aim of this study was to explore the effects of propofol on neural dynamics and EEG activity using the COALIA computational model, a whole-brain modeling framework that simulates brain-wide activity across multiple scales. By integrating both molecular and large-scale network-level effects, the model allowed us to investigate how propofol-induced changes in GABAergic transmission manifest in characteristic EEG rhythms and alterations in functional connectivity.

Our simulations successfully replicated several well-documented EEG features of propofol-induced anesthesia. As observed in the results, there was a marked increase in slow-wave (0.1–1 Hz) and alpha oscillations (8–12 Hz), particularly in the frontal cortical regions. These findings are consistent with the literature on the effects of propofol on brain activity during anesthesia, which has demonstrated that propofol enhances GABA_A receptor activity, prolonging IPSPs and leading to a reduction in overall cortical excitability (1, 3, 7).

The EEG signatures produced by the COALIA model under propofol anesthesia mirrored those found in clinical studies, particularly the emergence of large-amplitude slow-wave oscillations and synchronized alpha activity, which are considered markers of unconsciousness (5, 11). This coherence in the frontal cortices aligns with the breakdown of long-range thalamo-cortical and cortico-cortical communication under propofol, as shown by both our model and previous empirical findings (6).

Additionally, the model was able to simulate the transition from consciousness to unconsciousness, capturing the paradoxical excitation phase observed in clinical settings where beta oscillations (13–25 Hz) emerge before loss of consciousness. This provides further evidence for the robustness of the COALIA model in reproducing the temporal and spectral dynamics of anesthesia-induced brain activity (2, 8).

The use of a neural mass model for each cortical region and the inclusion of thalamic populations proved to be crucial in generating realistic EEG signals, as the thalamo-cortical network plays a central role in modulating alpha rhythms under anesthesia (9). The reciprocal interactions between the thalamus and cortex in the model created the sustained alpha activity and high spatial coherence across cortical regions, which is a key feature of propofol anesthesia (10).

Furthermore, the forward model, which projects the local field potentials (LFPs) onto the scalp, allowed us to generate simulated EEG data that closely matched empirical EEG recordings from human subjects undergoing propofol anesthesia. The power distribution of the simulated EEG, particularly the dominance of delta and alpha bands, further confirmed the model’s accuracy (Figures 3, 4, 5).

### Limitations and Future Work

While the COALIA model provides significant insights into the neural mechanisms underlying propofol anesthesia, there are several limitations to this study. First, the model assumes homogeneous activity within each cortical region, which may overlook more nuanced local variations in cortical excitability and dynamics. Additionally, the model relies on a fixed structural connectivity matrix derived from DTI data, which does not account for potential changes in connectivity that could occur dynamically under different levels of anesthesia.

Future work could address these limitations by incorporating finer-grained cortical models, allowing for more detailed simulations of local cortical dynamics. Another avenue for improvement would be to explore the impact of other anesthetic agents on brain activity to validate whether the COALIA model can generalize across different types of anesthesia. This could lead to a more comprehensive understanding of how different anesthetics modulate consciousness through similar or distinct mechanisms.

### Conclusion

In conclusion, the COALIA model provides a powerful framework for simulating the neural and EEG dynamics of propofol-induced anesthesia. By integrating molecular-level effects, such as the enhancement of GABA_A receptor activity, with large-scale network connectivity, the model replicates the characteristic slow-wave and alpha oscillations observed during anesthesia. The ability of the model to generate realistic EEG data that matches clinical recordings demonstrates its potential as a valuable tool for studying the neural mechanisms of consciousness and its modulation by anesthetic agents.

Through the simulation of propofol’s effects, this study contributes to a better understanding of the complex interactions between molecular mechanisms and large-scale brain networks, providing a foundation for future research on the neural correlates of consciousness and anesthesia.

## References

[1] John, E. R., Prichep, L. S., Kox, W., Valdes-Sosa, P., Bosch-Bayard, J., Aubert, E., … & Gugino, L. D. (2001). Invariant reversible QEEG effects of anesthetics. Consciousness and Cognition, 10(2), 165–183.

[2] Feshchenko, V. A., Veselis, R. A., & Reinsel, R. A. (2004). Propofol-induced alpha rhythm. Neuropsychobiology, 50(3), 257–266.

[3] Hales, T. G., & Lambert, J. J. (1991). The actions of propofol on inhibitory amino acid receptors. British Journal of Pharmacology, 104(3), 619–628.

[4] Kitamura, A., Marszalec, W., Yeh, J. Z., & Narahashi, T. (2003). Effects of halothane and propofol on excitatory and inhibitory synaptic transmission. Journal of Pharmacology and Experimental Therapeutics, 304(1), 162–171.

[5] Mhuircheartaigh, R. N., Warnaby, C. E., Rogers, R., Jbabdi, S., & Tracey, I. (2013). Slow-wave activity saturation and thalamocortical isolation during propofol anesthesia. Science Translational Medicine, 5(208), 208ra148.

[6] Casali, A. G., Gosseries, O., Rosanova, M., Boly, M., Sarasso, S., Casali, K. R., … & Massimini, M. (2013). A theoretically based index of consciousness independent of sensory processing and behavior. Science Translational Medicine, 5(198), 198ra105.

[7] Bhattacharya, B. S., Coyle, D., & Maguire, L. P. (2011). A thalamo– cortico–thalamic neural mass model to study alpha rhythms. Neural Networks, 24(6), 631–645.

[8] Breakspear, M. (2017). Dynamic models of large-scale brain activity. Nature Neuroscience, 20, 340–352.

[9] Esser, S. K., Hill, S. L., & Tononi, G. (2009). Breakdown of effective connectivity during slow wave sleep: investigating the mechanism of conscious disconnection. Journal of Neurophysiology, 102(4), 2096–2111.

[10] Modolo, J., Hassan, M., Deco, G., & Lefebvre, J. (2018). Probing the circuits of conscious perception with magnetophosphenes. bioRxiv. 10.1101/449769

[11] Dehaene, S., & Changeux, J. P. (2011). Experimental and theoretical approaches to conscious processing. Neuron, 70(2), 200–227.

[12] Tononi, G., & Edelman, G. M. (1998). Consciousness and complexity. Science, 282(5395), 1846–1851.

[13] Hudetz, A. G. (2012). General anesthesia and human brain connectivity. Brain Connectivity, 2(6), 291–302.

[14] Gomez, F., Phillips, C., Soddu, A., Boly, M., Schrouff, J., Vanhaudenhuyse, A., & Laureys, S. (2013). Changes in effective connectivity by propofol sedation. PLoS ONE, 8(8), e71370.

[15] Avena-Koenigsberger, A., Misic, B., & Sporns, O. (2017). Communication dynamics in complex brain networks. Nature Reviews Neuroscience, 19(1), 17–33.

[16] Massimini, M., Ferrarelli, F., Sarasso, S., Pigorini, A., & Tononi, G. (2010). Cortical reactivity and effective connectivity during REM sleep in humans. Cognitive Neuroscience, 1(3), 176–183.

[17] Friston, K. J. (2011). Functional and effective connectivity: A review. Brain Connectivity, 1(1), 13–36.

[18] Desikan, R. S., Ségonne, F., Fischl, B., Quinn, B. T., Dickerson, B. C., Blacker, D., … & Killiany, R. J. (2006). An automated labeling system for subdividing the human cerebral cortex on MRI scans into gyral based regions of interest. Neuroimage, 31(3), 968–980.

[19] Hagmann, P., Cammoun, L., Gigandet, X., Meuli, R., Honey, C. J., Wedeen, V. J., & Sporns, O. (2008). Mapping the structural core of human cerebral cortex. PLoS Biology, 6(7), e159.

